# Local Cortical Activity of Distant Brain Areas can Phase-Lock to the Respiratory Rhythm in the Freely Behaving Rat

**DOI:** 10.1101/259457

**Authors:** Daniel Rojas-Líbano, Jonathan Wimmer del Solar, Marcelo Aguilar, Rodrigo Montefusco-Siegmund, Pedro E. Maldonado

**Affiliations:** Laboratorio de Neurociencia Cognitiva y Social, Facultad de Psicología, Universidad Diego Portales, Santiago, Chile.; Unidad de Investigación y Desarrollo, Hospital El Carmen de Maipú, Santiago, Chile.; Programa de Neurología, Facultad de Ciencias Médicas, Universidad de Santiago de Chile, Santiago, Chile.; Department of Bioengineering, University of California, San Diego. La Jolla, CA 92093, USA; Escuela de Kinesiología, Facultad de Medicina, Universidad Austral de Chile, Valdivia, Chile; Department of Neuroscience & Biomedical Neuroscience Institute, Universidad de Chile, Santiago, Chile.

**Keywords:** Respiratory rhythm, sniffing, whisking, free behavior, Local Field Potential

## Abstract

An important unresolved question about neural processing is the mechanism by which distant brain areas coordinate their activities and relate their local processing to global neural events. A potential candidate for the local-global integration are slow rhythms such as respiration, which is also linked to sensory exploration. In this article, we asked if there are modulations of local cortical processing which are time-locked to (peripheral) sensory-motor exploratory rhythms. We studied rats freely behaving on an elevated platform where they would display exploratory and rest behaviors. Concurrent with behavior, we monitored orofacial sampling rhythms (whisking and sniffing) and local field potentials (LFP) from olfactory bulb, dorsal hippocampus, primary motor cortex, primary somatosensory cortex and primary visual cortex. We defined exploration as simultaneous whisking and sniffing above 5 Hz and found that this activity peaked at about 8 Hz. We considered rest as the absence of whisking and sniffing, and in this case, mean respiration occurred at about 3 Hz. We found a consistent shift across all areas toward these rhythm peaks accompanying behavioral state changes. We also found, across areas, that LFP gamma (70-100 Hz) amplitude could phase-lock to the animal’s respiratory rhythm, a finding indicative of respiration-locked changes in local processing. The respiratory rhythm, although occurring at the same frequencies of hippocampal theta, was not spectrally coherent with it, implying a different oscillator. Our results are consistent with the notion of respiration as a binder or integrator of activity between distant brain regions.

## INTRODUCTION

During the last decade, several studies have shown that animals produce different motor actions specifically tuned to obtain useful sensory information from the environment (Curtis and Kleinfeld, 2009; Najemnik and Geisler, 2005; Verhagen et al., 2007). These findings are part of what has been called ‘active sensation’ or ‘active sensing’, and examples are everywhere, from finger and eye movements in primates to echolocation in bats, electric sensation in fishes and whisking and sniffing in rodents (Hoffman et al., 2013; Schroeder et al., 2010). Sensory and motor activities characteristically coordinate during exploratory behaviors, in which the animal constantly gauges the environment through sensor-associated movements to gather sensory information.

The underlying message from these studies is that the coordinated or coupled activity of sensory and motor subsystems is what enables the animal to display adaptive behavior, contingent to the ongoing external changes. Thus, sensory-motor coupling allows distinguishing stimuli-triggered from internal activity (Ito et al. 2011; Cavanaugh et al. 2016), manipulating sensory inputs and enhancing stimulus detection (Carey et al. 2009; Verhagen et al. 2007; Rojas-Líbano and Kay, 2012), improving learning of skilled movements (Arce-McShane et al. 2016), and estimating the positions of objects (Mehta et al. 2007; Curtis and Kleinfeld, 2009). Accordingly, researchers have found a variety of activity correlations between the motor and sensory brain areas. Neural activity in sensory brain areas co-varies with parameters of motor sampling behaviors (Fee et al. 1997; Courtiol et al. 2011; Esclassan et al. 2012; Shusterman et al. 2011; Cury and Uchida, 2010; Ito et al. 2011; Rojas-Líbano et al. 2014) and activity in motor areas show stimuli-related features (Kleinfeld et al. 2002; Arce-McShane et al. 2014; Reimer and Hatsopoulos, 2010; Huber et al. 2012).

How do the activities of these exploratory sensory-motor processes relate to the local activity of the different brain areas? Depending on the contingencies of the situation, animals usually alternate between states of active exploration and periods of quietness or rest. Presumably, this implies an engagement and disengagement of a host of neural resources related to the control of motor sampling behaviors, to the processing of incoming signals, and to the coordination of these two. Given that is the entire animal that initiates and terminates behaviors such as exploration, we hypothesize that distant brain subsystems must also engage/disengage across these states. Therefore, an initial step is to evaluate if local processing in distant brain areas is related to the timing of the sensory-motor sampling rhythms, as well as to assess whether there are differences between exploratory behavior and rest or quiet states. In fact, the rat respiratory rhythm has been proposed as a sort of ‘master clock’ that can serve the role of coordinating the activity of diverse cerebral centers (Kleinfeld et al., 2014).

Here, we examined whether the local activity across distant brain areas can co-vary with the animal’s respiratory rhythm. We examined local field potential (LFP) activity across several brain areas in awake, freely behaving rats. We placed rats on a small elevated platform, where they naturally displayed both exploratory and resting states, and we did not deliver any specific sensory stimulation. Exploratory behavior was characterized by sniffing and whisking at about 8 Hz. During exploration, primary somatosensory (S1), primary motor (M1), primary visual (V1) cortical areas and dorsal hippocampus also exhibited regular activity at about 8 Hz. In contrast, at rest, the respiratory rhythm dropped to ~2.5 Hz, whisking lacked rhythmicity and S1, M1, V1 and hippocampus all switched to a common 5-6 Hz activity. In addition, we found modulations of gamma activity (70-100 Hz) in all neocortical areas, which were phase-locked to the animal’s respiratory rhythm. Our results suggest that low-frequency oscillations in the brain could serve the role of integration of activity across areas.

## MATERIALS AND METHODS

### Subjects and behavioral paradigm

Six adult male rats (three Wistar, three Sprague-Dawley, all 250-350 g), obtained from our animal facility (Bioterio Facultad de Medicina, Universidad de Chile), were used in the experiments. All rats were housed individually in standard clear polycarbonate home cages and maintained on a 14/10 h light/dark cycle (lights on at 8:00 P.M.). All recording sessions were carried out during the rats’ dark phase. All experiments followed institutional (protocol CBA0434 FMUCH) and NIH guidelines for the care and use of laboratory animals. We used an acrylic-made platform, with a 15 15 cm square surface, with a height of 25 cm. No objects were located nearby, no specific stimuli were delivered, and the platform was located at the center of a 45 cm x 70 cm x 70 cm wood-enclosed Faraday cage. The rats usually moved around the platform, whisking and perching at its edge without falling from it. Recording sessions lasted 10-15 min, with a total of four recording sessions from each rat. Sessions took place in complete darkness, and we monitored the rat both visually and through a video-camera under infrared illumination.

### Electrode fabrication

Electrodes for LFP recordings were made using 127 μm-diameter, polyimide-coated Nickel-Chrome wire (Kanthal, Palm Coast, FL), and were arranged in bundles of three leads per implantation site. On each bundle, three pieces of wire of ~2-cm long were glued together longitudinally, with a final inter-tip distance of ~500 μm. The impedance of individual LFP leads was kept in the range 50 - 100 kΩ at 1 kHz. Electromyogram (EMG) electrodes were made using 50 μm-diameter, PFA-coated tungsten wire (no. 795500, A-M Systems, Carlsborg, WA), according to the protocol reported by Berg and Kleinfeld (2003).

### Surgery

For all surgical procedures, we followed rodent aseptic surgery guidelines (Cunliffe-Beamer, 1993). We anesthetized the rat with isoflurane in a plastic induction chamber and then transferred it to a stereotaxic apparatus for the rest of the surgery. Anesthesia was maintained during surgery using isoflurane delivered at a modified nose cone in the stereotaxic apparatus. We monitored capillary oxygen saturation, heart rate, and respiratory rate during the entire surgical process. To implant EMG electrodes in the rat’s mystacial pad, we followed a previously published protocol (Berg and Kleinfeld 2003). We performed an incision along the head midline to expose the skull and opened subcutaneous pockets reaching the region posterior to the mystacial pad. We loaded a 25-gauge needle with three EMG leads, and inserted it through the subcutaneous pockets, posterior to the mystacial pad. The needle exited just anterior to the pad and was then removed, leaving the EMG leads sticking out of the skin. The leads were gently pulled back until they were within the pad, subcutaneously. Three leads were implanted on each pad, left and right. Two more leads were placed subcutaneously, dorsal to the nasal bone, to be used as a reference.

Craniotomies, 1-2 mm in diameter, were made to insert the electrodes. Locations were determined stereotaxically for the olfactory bulb (OB) (1.5 mm lateral from midline, 8.5 mm anterior to Bregma), whisker primary motor cortex (M1) (1.5 mm lateral from midline, 2 mm anterior to Bregma), whisker primary somatosensory (‘barrel’) cortex (S1) (5 mm lateral from midline, 2.7 mm posterior to Bregma) and reference (1 mm lateral from midline, 9 mm posterior to Bregma). In three of the six rats, we also implanted electrodes at dorsal hippocampus (2.2 mm lateral from midline, 3.8 mm posterior to Bregma) and primary visual cortex (V1) (2.4 mm lateral from midline, 8 mm posterior to Bregma). In OB, S1, M1 and V1, the tip of the electrode was gently lowered down to 1-1.5 mm from the brain’s surface. The hippocampus electrode was lowered to 3 mm. Electrode bundles in OB, M1, and S1 were implanted bilaterally. V1 and hippocampal electrode bundles were implanted only in the right hemisphere. For the reference, the electrode was placed right above the brain’s surface. Postsurgical infection prevention and analgesia were provided via daily intramuscular injections of ketoprofen and enrofloxacin 5% (Baytril^®^, Bayer), for seven days. Recording sessions were performed ten days after surgery at the earliest, depending on the rat’s condition.

### Chronic Electrophysiological Recordings

During recording sessions, signals were acquired using a head stage preamplifier (MPA 32I, Multichannel Systems, Reutlingen, Germany) providing the initial 10× amplification, connected to a Programmable Gain Amplifier (PG-32, Multichannel Systems). All channels were differentially amplified relative to the brain reference leads, wide-band recorded (1-5000 Hz) and digitized at 10 kHz using a PCI board (National Instruments, Austin, TX, USA). Custom-made scripts were written in LabWindows™/CVI software (National Instruments) for acquisition and on-line signal display.

### Data Analysis

Off-line analysis was performed using MATLAB ^®^ software (Release 2011b; The MathWorks, Natick, MA, USA). Custom code was written to import and process the data. We used functions from the Chronux toolbox (http://chronux.org), the Fieldtrip toolbox (Oostenveld et al., 2011), and custom-built functions. All EMG signals and nasal bone signals were first numerically high-pass filtered (100-1000 Hz) to remove any low-frequency artifacts, using a finite-response Equiripple Highpass filter designed using the MATLAB *firpm* function. The difference of two raw EMG channels formed the bipolar EMG. Each bipolar EMG was rectified and smoothed using a moving-mean filter with a 30-ms window, to obtain the final rectified-and-smoothed EMG (rsEMG), which was used for spectral analysis and time-domain whisking frequency calculations. The use of rsEMG, or ‘integrated’ EMG, is customary in these analyses (Berg and Kleinfeld, 2003; Rojas-Líbano et al., 2012; Rangayyan, 2002; Strittmatter and Schadt, 2007).

#### Behavioral classification

The signals from the OB and the whisker pad were used to assign each time point of the session to a behavioral state. We used the low-passed (below 15 Hz) OB LFP as a proxy for the rat’s respiration, as previously shown (Courtiol et al. 2011; Carlson et al. 2014; Jessberger et al. 2016; Nguyen Chi et al. 2016; Rojas-Líbano et al., 2014). Following Nguyen Chi et al., 2016, we refer to this signal as the respiratory rhythm (RR). Using an automated algorithm, we detected periods in which the rat was whisking (W+), not whisking (W−) (from the whisker EMG signal), and sniffing (S+), or not sniffing (S−) (from the OB LFP signal). Hence, we assigned each time point into one out of four categories (W−/S−; W−/S+; W+/S−; W+/S+). Sniffing categorization was performed by obtaining the peak frequency in each time window of the olfactory bulb signal spectrogram. The spectrogram was obtained for frequencies between 1 and 12 Hz using windows of 1 s with a 0.1 s overlay, using three tapers for a bandwidth of 2 Hz. To obtain a clear peak frequency, the spectrum of each time window was smoothed though a moving mean with a window of the calculated bandwidth. The peak frequency was then selected. For those time windows where no clear peak or more than one peak was obtained, the value from the previous time window was kept. This procedure was performed forward and backward in time to avoid phase distortions. When classification performed in both directions was not the same, interpolation of the last and first next coincident frequency was accounted as the peak frequency. When the peak frequency was above 6 Hz the period was classified as sniffing (S+), if it was below 5 Hz it was classified as not sniffing (S−). The performance of the classifier algorithm agreed quite well with visual inspection of the data.

For whisking classification, the variability of the whiskers’ rsEMG signal in time domain was assessed for each session. The moving standard deviation of the signal calculated with a 0.5-s window was obtained. The minimum standard deviation obtained was used to classify the signal. Those periods in which the standard deviation was greater than eight times the minimum standard deviation of that session were classified as whisking periods (W+), otherwise were classified as not whisking periods (W−). The automatic classifier accuracy was assessed visually, comparing the results with a manual categorization performed by a trained experimenter looking at the signal in the time and frequency domain. For the calculation of spectra per area per behavioral state, the time periods classified according to each behavioral condition were subdivided into 3-seconds epochs to be used as trials. Spectral estimates were obtained using multitaper spectral analysis.

#### Peri-respiratory activity across brain areas

We also sorted peri-respiratory LFP epochs for each brain area to compare across behavioral states in the following way. First, we filtered the OB LFP below 15 Hz and searched for local minima of the time series to determine the LFP RR cycle onsets. After collecting all the cycles, we parsed them in two groups, according to their cycle frequency: below 5 Hz and above 6 Hz (cycles with frequencies in between 5 and 6 Hz were not used for this analysis). These corresponded to our previously defined Rest and Exploration states, respectively (because sniffing without whisking was very rare; see Results). We then used the timestamps of each OB cycle onset to collect the corresponding peri-respiration epochs from all the channels. In this way, we were able to construct two matrices of LFP epochs from each brain area, one per behavioral state. Examples of these data for one rat are shown in figure 4.

The matrices of peri-respiration epochs were also used to compute the spectral coherence between the RR and the gamma activity from each brain area. We filtered the peri-respiration epochs to obtain a gamma (70-100 Hz) signal and then calculated its envelope through the absolute value of the Hilbert transform of the filtered data. We then computed the multi-taper spectral coherence between the gamma envelope of each epoch and the corresponding respiratory waveform (2 tapers, bandwidth W=2 Hz). We did this for all epochs from each brain area and rat. To construct a null model, we randomly permuted (1000 times) the gamma envelope – respiratory wave pairs, within each area and rat, and computed the coherence from this new (randomized) data set. We then computed, for each frequency point, the 95^th^ percentile of the randomized data set, and used it as a significance threshold (see Figure 5). Next, we defined a frequency range [−1.5, 1.5] Hz centered at the spectral maximum of the RR of each rat. We then counted the number of statistically significant points within this frequency range (see Figure 5).

#### Current source density

The three voltage time series recorded from each brain location were used to compute the current-source density (CSD) in each area, which corresponds to the second spatial derivative of the potential on depth (Nicholson and Freeman 1975). CSD was calculated separately from each area as CSD = −[LFP_1_(t) − 2·LFP_2_(t) + LFP_3_(t)]/Δz^2^, where LFP_*i*_ is the voltage recorded from electrode *i* and Δz corresponds to the vertical separation between electrodes, i.e., Δz = 500 μm.

#### Cluster-based statistics

To compare between power spectra from different behavioral states, we implemented non-parametric, cluster-corrected statistical comparisons (Maris and Oostenveld, 2007). In our case, instead of trials, we had the power spectra calculated from the 2-second epochs assigned to either rest or exploration states. First, for every sample (i.e., each (power, frequency) value) we compared the two spectra using a t-statistic, obtaining the non-permuted t-values. We then permuted the epochs, assigning them randomly to each of two groups, and calculated again the t-values. We repeated the permutation procedure 1000 times, obtaining a distribution of t-values for each sample. We then selected all the samples with non-permuted t-values corresponding to p < 0.05 of the distribution. Finally, we performed the cluster-based correction of the t-values. To do this, we clustered the selected samples by adjacency in the frequency axis and calculated cluster-level statistics by taking the sum of the t-values within a cluster.

## RESULTS

We used a training-free paradigm, in which rats were placed on an elevated platform, to specifically avoid repetitive stereotyped behaviors characteristic of training in conditioning tasks. It is known that rats usually perch on the edge of elevated platforms and can be trained to whisk in the air to get a reward (Fee et al. 1997; Ganguly and Kleinfeld 2004). Without giving any rewards, we found that rats regularly displayed quiet resting and exploration periods, as assessed visually and by whisking and sniffing recordings (see Figs. 1 and 2). Rats usually alternated between periods of active exploring and quietness or minimal overt activity. In all the rats (n = 6) we recorded the LFP activity of the olfactory bulb (OB), the whisker primary somatosensory cortex (S1), the whisker primary motor cortex (M1), and the EMG activity of the whisker pads. In a subset (n = 3) of the rats, we additionally monitored the LFP activity in primary visual cortex (V1) and dorsal hippocampus (Hipp).

**Figure 1.**
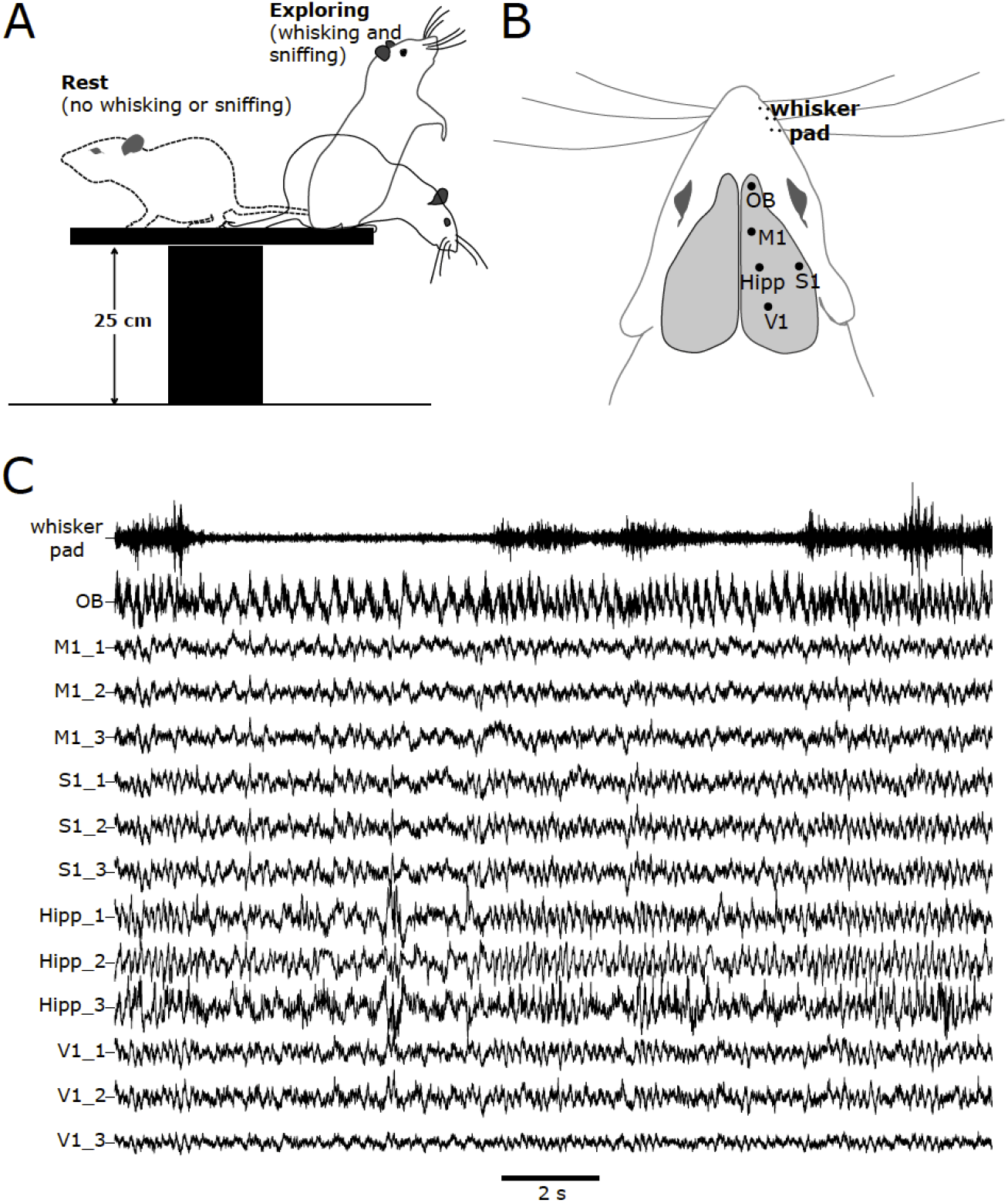
Behavioral setup, areas recorded and raw time series examples. *A*: Schematics of the elevated platform and rat behaviors. *B*: Schematic drawing of a rat’s head viewed from above indicating recording sites: whisker pad, where we placed subcutaneous electrodes, and different cerebral locations, where we implanted chronic electrodes. *C*: Example traces of LFP time series from each brain area recorded, and the EMG of the whisker pad. Note the coincidence of periods of slow oscillations (about 2-3 Hz) in the OB with periods of no whisker EMG activity, and the coincidence of periods of 6-8 Hz activity in the OB with periods of whisker EMG activity. Three probes vertically separated by 500 μm were implanted in each brain area, with the top probe (#1) located on the brain’s surface. All brain channels were recorded at the same amplification and had the same reference. Drawings and data are shown only in one hemisphere for purposes of clarity. Data are from Rat 4.

**Figure 2.**
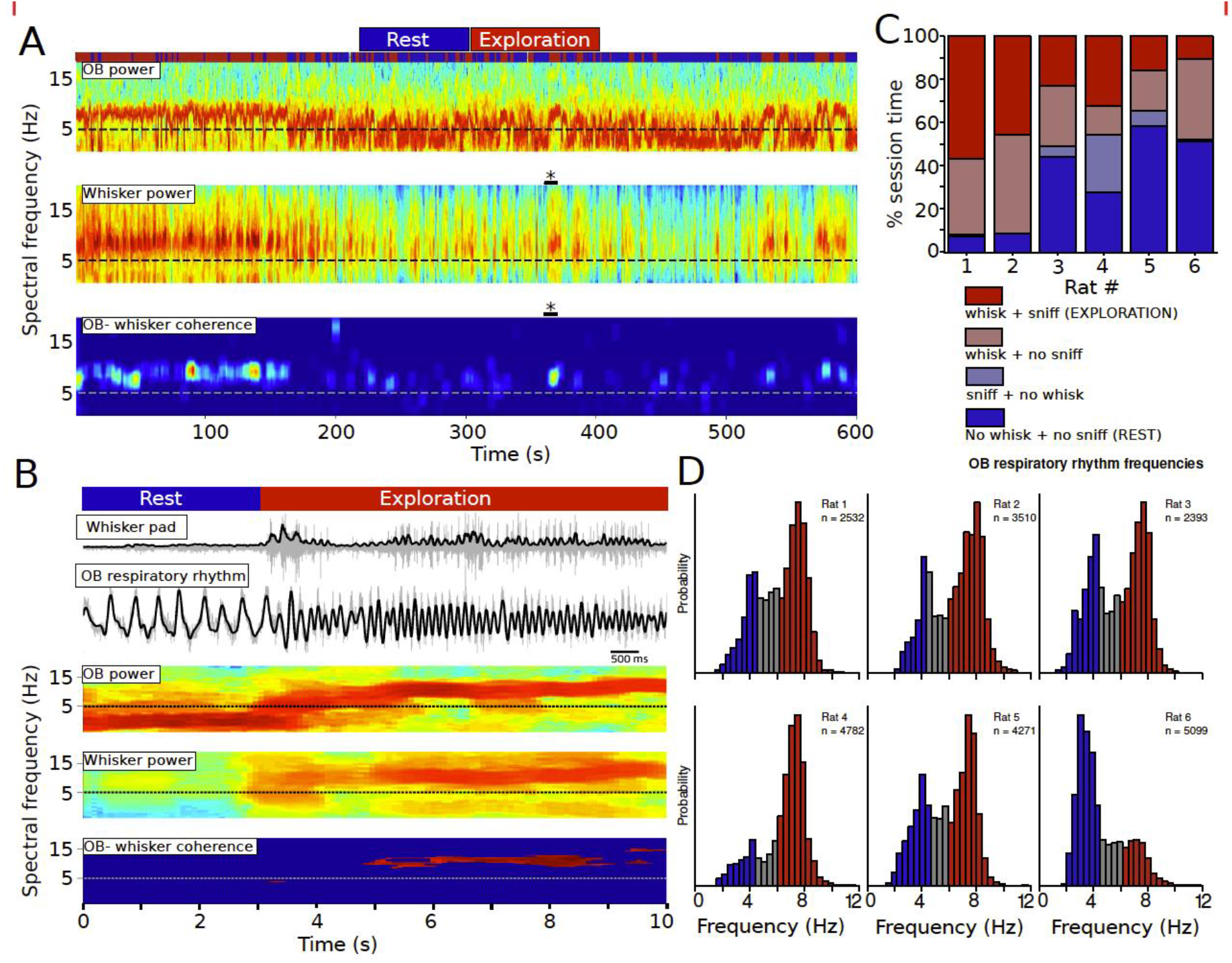
OB LFP and whisker EMG spectral frequencies during a recording session, and distribution of the two behavioral states. *A*: The top plot shows a spectrogram of the raw OB LFP for a 10-minute session, displaying frequencies from 0 to 20 Hz. Largest power values (dark red colors in the spectrogram) indicate the respiratory/sniffing frequency of the rat. The middle plot shows the corresponding spectrogram for the rectified and smoothed electromyogram (rsEMG) of the whisker muscles. The bottom plot shows the spectral coherence calculated for the two signals. Only significant values of coherence are plotted in the color scale (non-significant values are colored dark blue). Coherence occurs for the most part above 5 Hz. The timepoints of the entire session were classified as ‘Rest’ or ‘Exploration’ based on the co-occurrence of whisking and sniffing (see the two-colored line at the top of the figure). We drew a line at 5 Hz to aid the eye. Data are from Rat 1. *B*: Example of a transition between Rest and Exploration states. The data correspond to an enlargement of the section marked with a black bar and an asterisk in A. Top: the two raw (gray) and filtered (black) signals we used for behavior classification, whisker EMG and OB LFP. The Rest state was characterized by low-frequency (1 - 3 Hz), high-amplitude OB activity and no muscular whisker activity (flat EMG), whereas the Exploration state showed high frequency (above 5 Hz), low-amplitude OB activity and whisker EMG activity. The spectrograms of both signals and the dynamic spectral coherence between them are shown. Note augmented coherence during the Exploration period. A line was drawn at 5 Hz to aid the eye. *C*: Relative frequency of the different behaviors, expressed as the percentage of session time for each rat. *D*: Histograms of OB respiratory rhythm frequencies, with all sessions pooled for each rat. In all plots, dark red corresponds to ‘Exploration’ behavior and dark blue to ‘Rest’ behavior.

We used the low-passed (below 15 Hz) OB LFP as a proxy for the rat’s RR, and the whisker pad EMG as a measure of whisking. Using whisking and sniffing as indicators of rest and exploration states, we could parse periods throughout the session and analyze the main underlying LFP rhythms and their relationships with the RR. Figure 1C shows an example of all the LFPs and the EMG recorded simultaneously in one rat.

### Behavior analysis

On each recording session, each rat (n = 6) was placed on top of a small elevated platform (see Materials and Methods), with no objects within reach, and without experimental delivery of any particular sensory stimulus. Typically, rat behavior was characterized by alternating periods of active exploration with rest (see Figs. 1 and 2). Exploratory periods included displacement within the platform, up and down head movements, and typically, positioning at the edge of the platform displaying vigorous whisking and sniffing, orienting their nose toward outside the platform. In contrast with the exploration state, rats also displayed periods of rest, in which they showed no overt signs of activity besides breathing and occasional small movements.

Using an off-line algorithm (see Methods) we classified each time point as sniffing/no-sniffing and also as whisking/no-whisking. Therefore, we were able to parse the entire session time in the four possible combinations of these two behaviors (see Fig. 2). We found a range of 10-50% of session time for simultaneous whisking and sniffing, and a range of 7-58% of session time for the simultaneous absence of whisking and sniffing. Regarding the remnant categories, we found that sniffing without whisking was relatively rare (range 0.1-26%), except for one rat, and that whisking without sniffing, was more common (13-45 %).

Whisking and sniffing, which we defined as activity above 6 Hz, usually peaked between 7-9 Hz (see Figs. 1 and 2). The simultaneous presence of the two sampling modes indicates a behavior that is known to be centrally coordinated (Moore et al., 2013: Kleinfeld et al., 2014). Therefore, we used this sniffing/whisking simultaneity as our marker of an active behavioral state, which we refer to as the ‘Exploration’ state throughout the text. The simultaneous absence of sniffing and whisking was our marker of a more passive behavioral state, which we refer to as the ‘Rest’ state throughout the text.

These results show that in our experimental setup, rats displayed spontaneously different behaviors such as rest and active exploration. Our neural and muscle recordings allowed us to monitor the sensory-motor activity underpinning these behaviors and hence to analyze the activity across brain areas in these two states.

### Local activity across areas in the frequency range of the sensory-motor exploratory rhythms (below 15 Hz)

We evaluated the coordination between distant brain areas during exploration and rest behaviors. Initially, we focused on lower frequencies (below 15 Hz), a range of frequencies in which sensory-motor exploratory behaviors occur. We sought to determine whether active sensing behavior and neural activity across distant brain areas displayed common oscillatory frequencies. In Figure 3 we show the power spectra below 15 Hz of all recorded areas, averaged over animals and parsed by behavioral state. We used these spectra to search for consistent frequency shifts across behavioral states at all areas recorded.

**Figure 3.**
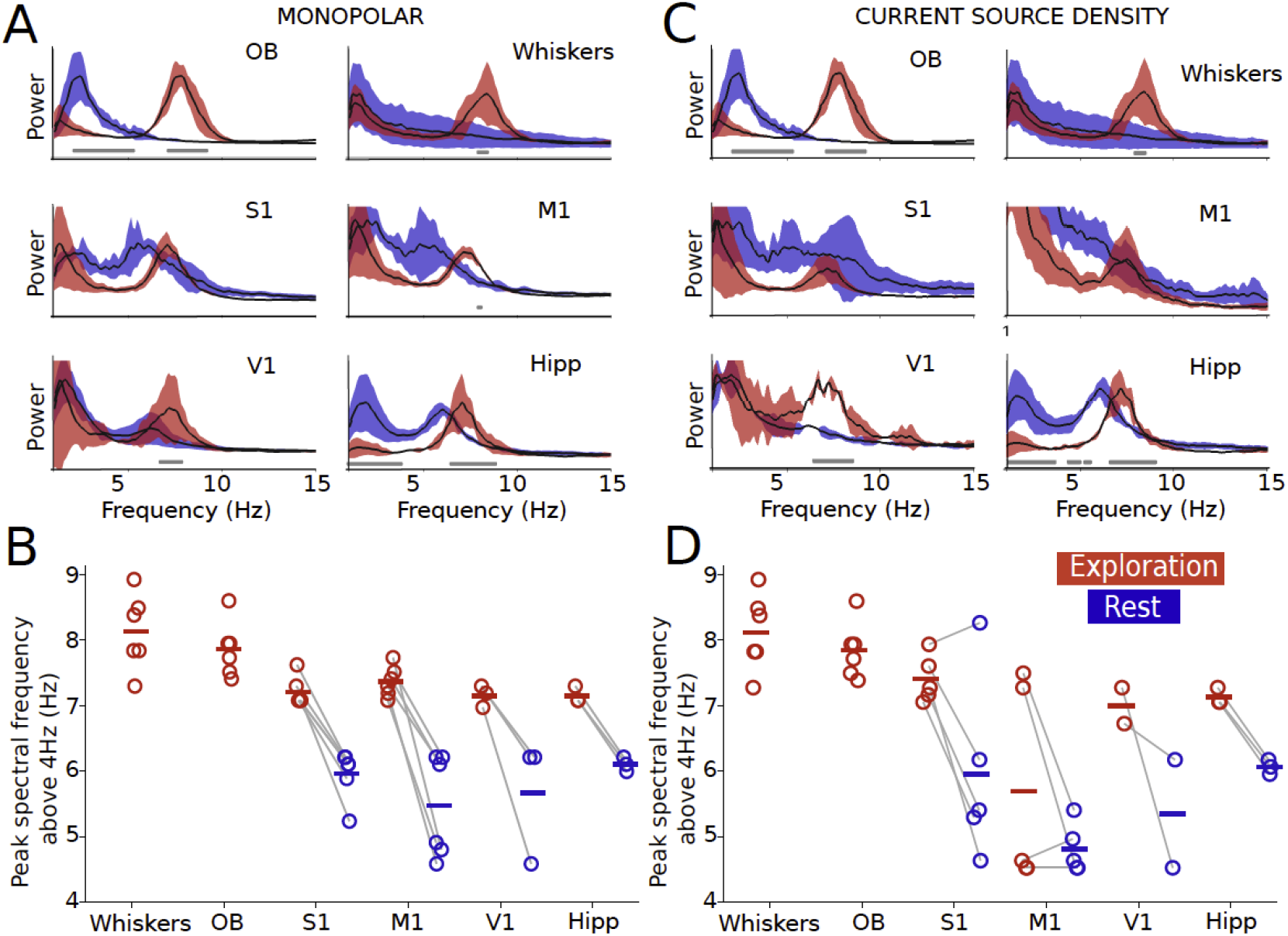
Average power spectra of the different areas, parsed by behavioral state. In *A* and *B*, power spectra were calculated using the raw signal, which was recorded in a monopolar derivation (i.e., all areas referenced to a common distant brain site). In *C* and *D*, power spectra were calculated using the CSD, which was calculated using the time series from the three leads implanted in each area. *A*: Each spectrum is the average across all the rats (n = 6 for OB, whiskers, M1, S1; n = 3 for V1, Hipp). Note that in the Exploration state both state indicators (OB and whiskers) display a peak of activity around 8 Hz, and similar peaks are seen in the rest of the brain areas. In the Rest state the peaks of activity are shifted to a lower frequency, around 6 Hz, again consistently across animals and brain areas. *B*: Power spectra maxima (above 4 Hz) for each rat, parsed by behavioral state. This plot shows detailed per-rat information from the spectra averages presented in *A*. Each rat has two peak values, one for Exploration and other for Rest. Note the consistent change in frequency that occurs in each animal with the change in behavioral state. Values in both states are linked by grey lines for each rat. *C*: Using the CSD, spectra were similar to the monopolar case. It is noisier in V1 due to the use of only two rats. CSD were only calculated for brain signals and therefore whisker EMG spectra are the same. *D*: CSD spectral peaks also showed a tendency to decrease from exploration to rest. One rat in S1 and three rats in M1 did not followed this pattern. In the case of M1, this happened because the spectra did not contain clear peaks above 4 Hz.

For the following statistical description (means and standard deviations), we used n = 6 rats for OB, EMG, S1, and n = 3 rats for V1 and Hipp. In the Rest state, cortical areas showed clear oscillation peaks closer to hippocampal theta activity (See Table 1). As defined for this state, there was no sniffing and no whisking. On the contrary, when we detected that the animal had turned to the exploration state, the average sniffing frequency was 7.8 ± 0.5 Hz and the average whisking frequency was 8.1 ± 0.7 Hz. In this state, cortical areas shifted their power maxima closer to the whisking and sniffing frequency, although at a slightly lower frequency (see Table 1). This shift from about 6 to 7 Hz that occurred in cortical areas and hippocampus was consistent across animals (see Fig. 3B). These results demonstrate that different motor and sensory areas can display similar oscillatory patterns that shift coherently with behavior.

**Table 1.** Means and standard deviations of power peaks across brain areas in the 4-15 Hz range, for Rest and Exploration conditions. We calculated the power spectra from each area and rat and then produced a mean spectrum across rats. We did this using the raw (i.e. monopolar) recordings and using the CSD computed from the three leads on each area. For OB, M1 and S1, n = 6 rats. For V1 and Hipp, n = 3 rats. OB activity peaked around 2-3 Hz during rest behavior.

|  |  | OB | M1 | S1 | V1 | Hipp |
| --- | --- | --- | --- | --- | --- | --- |
| Monopolar | REST | - | $5.6 \pm 0.7$ | $5.9 \pm 0.4$ | $5.3 \pm 1.2$ | $6.1 \pm 0.1$ |
| | EXPLORATION | $7.8 \pm 0.5$ | $7.4 \pm 0.2$ | $7.2 \pm 0.2$ | $7.1 \pm 0.2$ | $7.2 \pm 0.2$ |
| CSD | REST | - | $4.7 \pm 0.4$ | $5.9 \pm 1.4$ | $5.3 \pm 1.2$ | $6.0 \pm 0.1$ |
| | EXPLORATION | $7.8 \pm 0.4$ | $5.7 \pm 1.6$ | $7.4 \pm 0.4$ | $7.0 \pm 0.4$ | $7.1 \pm 0.1$ |

Given that our recordings used a monopolar derivation, there exists the possibility of electrotonic spread of voltage from remote sources, which could account for the previous results. To control for this possibility, we computed the CSD from each brain area and used it to obtain again the spectral estimates. The CSD is a function of the three timeseries from the three leads implanted in each area, with the topmost lead located at the brain’s surface and with a vertical inter-lead separation of 500 μm. It is therefore a local signal that should not be affected by passive spread from remote oscillators, given that its computation involves a subtraction from nearby electrode leads (Bastos and Schoffelen, 2016). The results were similar, albeit noisier, to the monopolar case (see Figs. 3C, 3D and Table 1). Therefore, the rhythms we detected on each area, in both behavioral states, were likely of local origin.

These results show that frequencies of sensory-motor exploration were apparent in all the brain areas recorded. During exploration, we detected peaks across areas that were closer to the frequency of the sensory-motor exploratory rhythms, sniffing and whisking. In contrast, during rest those peaks moved to lower frequencies.

### Local activity across areas in the gamma frequency range (70-100 Hz)

To assess whether local activity in the recorded areas was related to the timing of the sensory-motor sampling rhythms, we investigated LFP frequencies above 70 Hz. Because we wanted to assess phase-locking, we needed a precise reference time point to align local activity. EMG recordings were not always artifact-free or had contraction onset and offset times that were not easy to detect on a cycle-by-cycle basis. Therefore, and considering that for the most part our rats did not display sniffing without whisking, we used only the OB RR cycles, which had clear onsets, to align the data from all the areas.

Using the time frame provided by the OB RR, we collected 2-second LFP epochs, centered at the RR onset, from all brain areas. Using all collected RR cycles from each rat (for numbers of cycles, see histograms in Fig. 2), we calculated respiration-locked spectrograms and assessed for modulations of activity above 70 Hz. An initial inspection of the spectrograms showed several examples of power fluctuations that were time-locked to the respiratory rhythm (see supplementary Fig. 1). To assess for the relation between gamma power changes and the respiratory rhythm, we first computed correlation functions in the time domain. We calculated the cross-correlations between the mean RR and the mean time-varying gamma power aligned to the RR onset (See supplementary Fig. 2). We compared this with a cross-correlation between the mean RR and gamma power fluctuations aligned to a matching amount of randomly selected points in time, uniformly distributed across the session. This analysis showed that gamma power across rats and brain areas was indeed fluctuating at the respiratory rhythm’s rate.

Given the previous result, which pointed to the modulation of local gamma activity by the RR, we then assessed the effect through a frequency domain analysis. Time-domain estimates suffer from bias arising from the fact that measures at different time points are not independent from each other (Jarvis and Mitra, 2001). We therefore computed the multi taper spectral coherence (2 tapers, bandwidth W = 2 Hz) between the envelope of the gamma-filtered LFP of each area and the rat’s RR (Fig. 5). We compared the actual coherence values with the coherence computed from a surrogate data set in which we permuted the RR and gamma envelope pairs. We used as a significance threshold the 95^th^ percentile of the permuted coherence values, calculated for each frequency point. Specifically, we assessed the coherence in the frequency range of the RR. We centered this range at the frequency with larger power in the RR spectrum, ±1.5 Hz. We display the data by computing the fraction of significantly different frequency points (i.e., frequency points at which the actual coherence value was larger than the coherence value of the 95^th^ percentile of the permuted data) from the RR range (see Fig. 5C). As assessed by this method, all rats showed some extent of gamma-RR coherence in some areas, indicating that local gamma can phase-lock to the animal’s RR. During exploration, M1 showed the largest fraction significant, followed by V1, Hipp and then S1. In average, the fraction of significant frequency points was larger for exploration (see Fig. 5C and Supplementary Fig. 3).

These analyses showed that in all the rats and brain areas monitored, the amplitude of local gamma activity could phase-lock to the animal’s respiratory activity. This local-global coherence was smaller for basal respiration than for exploration.

### The olfactory bulb and hippocampal rhythms during behavioral states

Given that the hippocampus produces theta activity, which displays a range of frequency similar to that of the exploratory rhythms, we investigated further the relation between hippocampal and respiratory rhythms. We found that the RR frequency range and descriptive statistics were quite consistent across rats, in both the exploration and rest states (see Fig. 2D and Fig. 3). Characteristically, in the OB the RR was coordinated with faster (above 40 Hz) oscillations. Consistently, each RR cycle contained one gamma burst, which started roughly in the middle of the RR cycle, as expected from previous reports (Rojas-Líbano and Kay, 2008). These gamma bursts started at frequencies about 70-100 Hz and progressively decreased to frequencies close to 40-50 Hz (see Fig. 6). During exploration, 70-100 Hz were much more prominent than 40-50 Hz, and the converse was true for the rest state, were the 40-50 Hz had a larger power, as previously shown (Kay, 2003).

Recordings from the hippocampus showed the characteristic theta activity, with spectral peaks around 6-8 Hz. However, they were not related in an obvious way to the OB RR (see Fig. 6). When analyzing spectrograms and LFP averages locked to the onset of the OB’s respiratory rhythm, hippocampal activity did not display the same consistent trend across rats. This was especially true in the exploration case. Time-locked spectra showed a plain 1/f profile, and time-locked average LFP were different across rats and of lesser amplitude. Nevertheless, during rest two of the three rats displayed a hippocampal respiration-triggered average which resembled that of the OB RR (see Fig. 6). Across the sessions, OB and Hippocampus clearly shared a frequency range, but their activity was rarely coherent (see Fig. 7). All three rats with hippocampal electrodes displayed the same relationship with OB. These results suggest that the OB respiratory rhythm and the hippocampal theta rhythm are, or derive from, two separate pacemakers. Because of the projections of OB and hippocampus, they both have the potential to modulate activity in cortical areas.

## DISCUSSION

We recorded local field potentials from several brain areas in freely behaving rats placed on top of a small platform. Our aim was to assess whether an endogenous rhythm such as respiration could phase-lock to local neural processing across distant areas. We used sensory-motor exploratory rhythms, specifically whisking and sniffing, to identify two behavioral states: rest and exploration.

In our setup rats spontaneously displayed orofacial exploratory behaviors, such as whisking, sniffing, and also periods of rest. We found whisking at 7-9 Hz (Figs. 2 and 3), a frequency range consistent with previous reports of free-ranging rats (Semba and Komisaruk, 1984; Fee et al., 1997; Berg and Kleinfeld, 2003, O’Connor et al., 2002). The OB LFP in the 1-15 Hz range provides an excellent correlate of the animal’s respiratory rhythm, in behaving rodents (Courtiol et al., 2011; Lockmann et al., 2016; Nguyen Chi et al., 2016; Jessberger et al., 2016; Rojas-Líbano et al., 2014). We used this measure to monitor the breathing rhythm. Previously reported sniffing frequencies in rats go from 6 to 10 Hz in freely moving conditions (Tsanov et al., 2014; Rojas-Líbano et al., 2014), consistent with our findings. Moreover, whisking and sniffing occurred simultaneously, and in a spectrally coherent way (Fig. 2), a behavioral signature of orofacial exploration (Moore et al., 2013).

During the exploratory behavior, all cortical areas showed spectral peaks of activity closer to whisking-sniffing rhythms (Fig. 3). Since monopolar extracellular recordings (in our case, all channels recorded with respect to a posterior location in the brain’s surface) like ours can potentially display activity passively spread to the recording electrode from remote oscillators, we also searched for proxies of intrinsic, local activity within each area. It has been shown that CSD measures from both cortex (Ahrens and Kleinfeld, 2004) and hippocampus (Berg et al., 2006) are relevant readouts of activity in awake rats. Therefore, we calculated spectra using the raw recordings and using the CSD time series from each area. Being a signal computed as a function of voltage time series from three closely spaced (500 μm in our case) electrodes, it is unlikely to contain remotely generated activity. CSD spectra also showed clear peak shifts mirroring exploration and rest behaviors (Fig. 3). CSD spectral peaks were more variable than the monopolar ones. Since our study analyzed activity simultaneously recorded across distant brain areas, spectral peaks were calculated first as a within-rat average over all periods, parsed by behavioral state, and then as a group average. Our averages across periods of exploration can potentially blur subtle local-global differences between periods. We interpret the differences between monopolar and CSD results as reflecting a balance between local, intra-area processing and incoming, more globally broadcasted signals (i.e., the sensory-motor exploration rhythms). The interplay between local activity and incoming rhythms probably depends on the specific contingencies of the situation, such as the degree of attention, which could add more weight to either local or global activity.

We then examined whether local (i.e., intra-area) neural processing across distant brain areas was related to the sensory-motor exploratory rhythms. We calculated the spectral power in the gamma range (70-100 Hz) of LFP epochs aligned to the onset of the respiratory rhythm, across all areas (Fig. 4). We indeed found significant changes in gamma power that were phase-locked to the respiratory rhythm in all areas recorded (Fig. 5 and Supplementary Figs. 1, 2, 3). Besides local processing phase-locked to the exploratory rhythms in orofacial-related areas, we found that V1 and hippocampus also displayed periods of gamma modulation. This suggests that exploratory rhythms can impact local processing across vastly separated brain areas, even outside the specific sensory system associated with the exploratory behavior. Presumably, this modulation would depend on time-varying situational requirements, both internal and external, which we did not control in our design. These variations probably explain the differences found across rats and areas in terms of modulation amplitude and occurrence. When comparing across behaviors, our data showed that phase-locking of gamma and respiration was more common during exploratory behavior than during rest (see Fig. 5C). This is similar to a recent report in humans (Herrero et al., 2018), where for the most part volitional breathing, and not basal breathing, displayed phase-locking of local gamma with the subject’s respiration.

**Figure 4.**
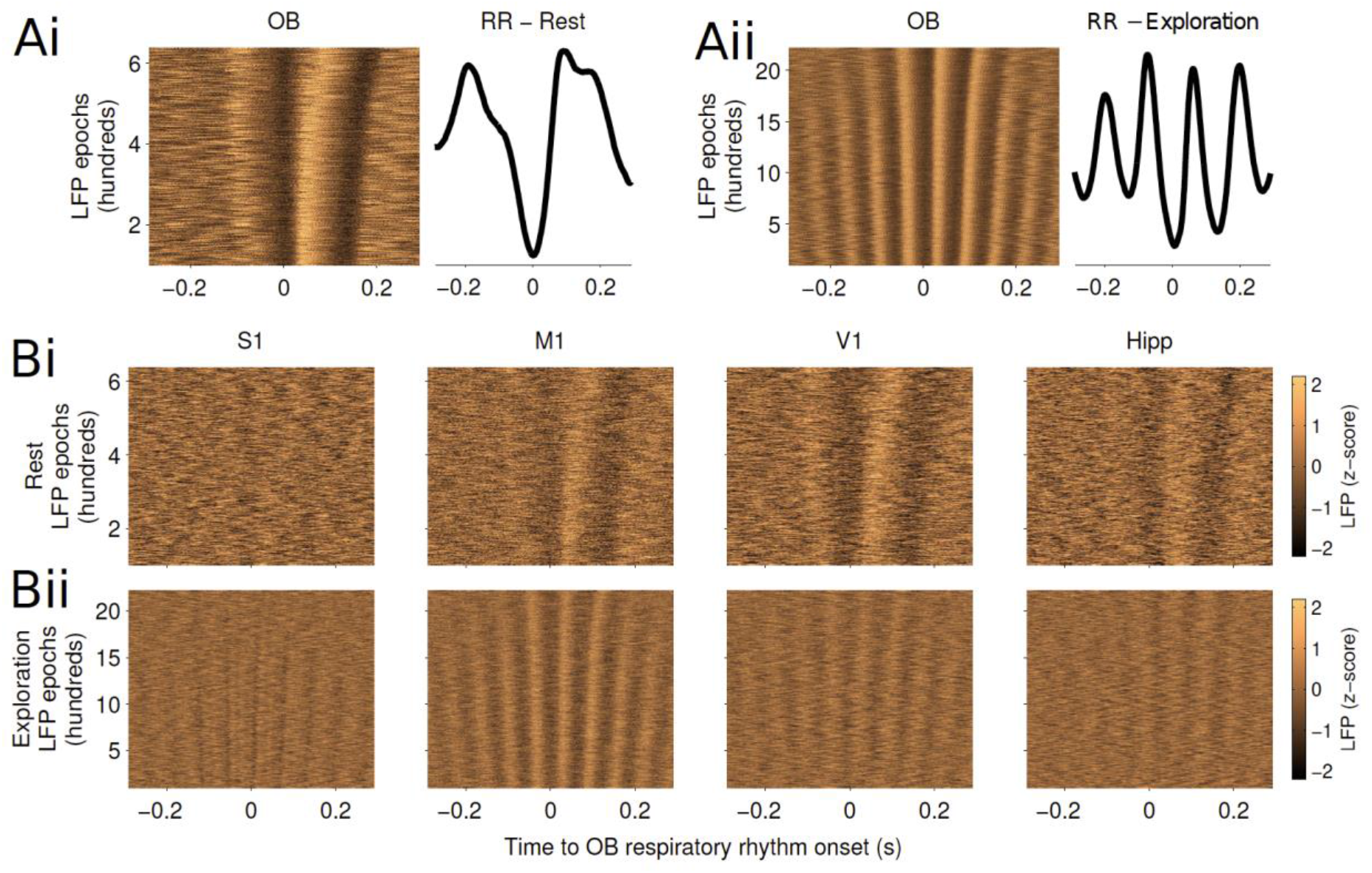
Examples of peri-respiration LFP epochs across brain areas and behavioral states. *A*: OB LFP epochs from which the mean RR waveform was calculated. *Ai*: Rest behavior. In the raster plot (left side), peri-respiration LFP epochs are shown. Epochs are centered at the start of the OB cycle, with dark tones corresponding to negative values and lighter tones to positive values. LFP data was z-scored separately for each epoch. The mean respiratory rhythm (RR, right side) correspond to the averaged OB LFP activity across epochs. *Aii*: Exploration behavior. Data organization as in *Ai*. *Bi*: Raster plots of peri-respiration LFP epochs for the different brain areas recorded, corresponding to the rest behavior. *Bii*: Exploration behavior. Data organization as in *Bi*. Data are from rat 5.

**Figure 5.**
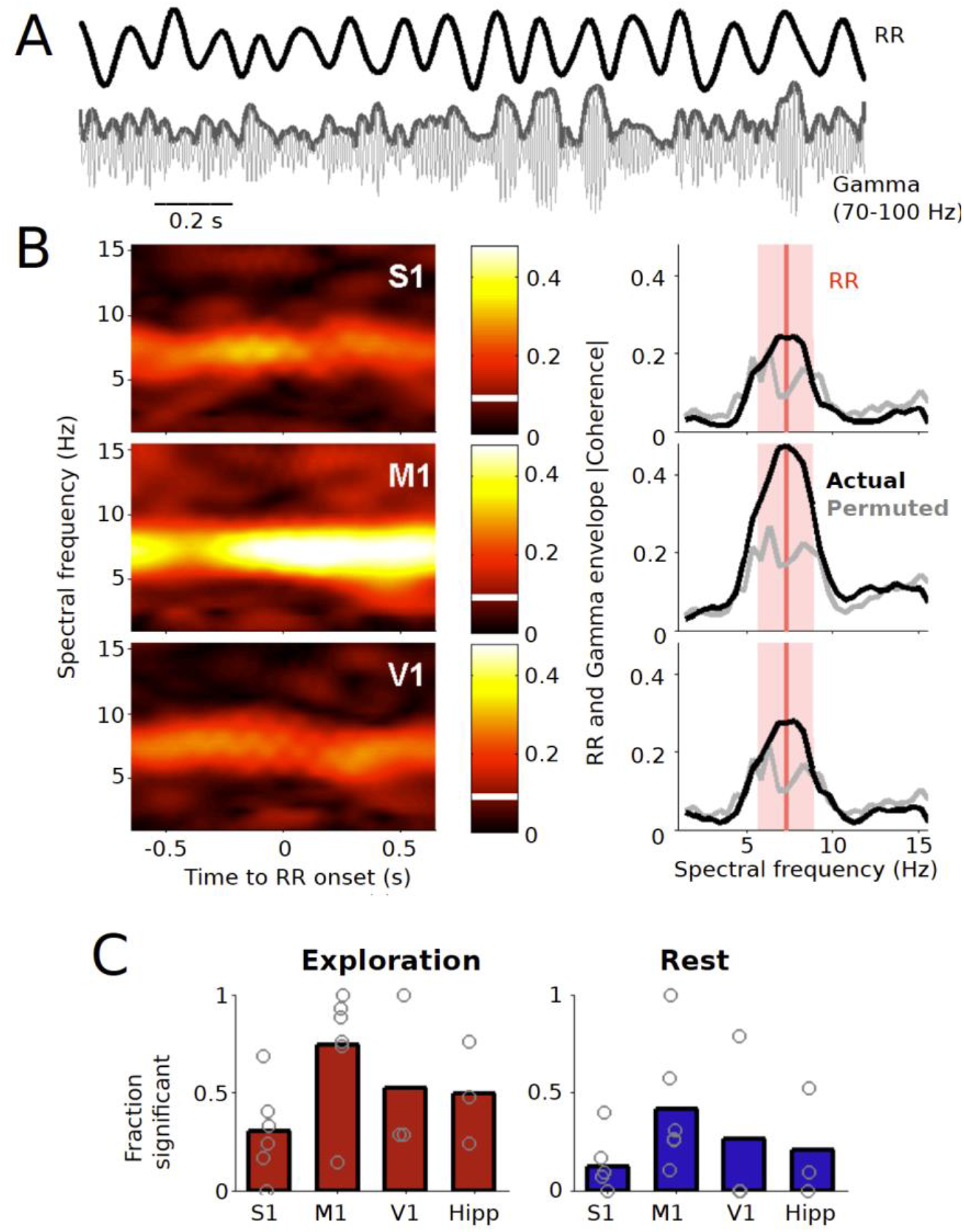
Spectral coherence between respiratory rhythm and gamma power. *A*: Example traces from the respiratory rhythm (RR) and the simultaneously recorded LFP activity from M1 during exploratory behavior, shown here as filtered data between 70 and 100 Hz. Thick grey line on top pf gamma represents the envelope, calculated from the absolute value of the Hilbert transform of the filtered time series. Data are from rat 5. *B*: Spectral coherence between the rat’s RR and the gamma envelope. Left-side plots show the coherencegrams centered at the time of RR onset, and right-side plots show the coherence calculated on the interval −0.5 to 0.5 seconds with respect to RR onset. Thick black line shows the absolute value of the coherence. Grey line represents the 95^th^ percentile of the coherence values calculated for the permuted (1000 times) data set. The red vertical line represents the peak value of the RR spectrum and the pale red area represents a range of ±1.5 Hz over which significance between actual and permuted coherence was assessed. *C*: Fraction of frequency points in the RR peak ±1.5 Hz range for which actual coherence was larger than the permuted values. Bars represent means over rats and dots represent mean values for each rat.

The rhythms of orofacial exploration occurred in the same range of activity as hippocampal LFP rhythms, which in our data displayed power maxima around 7 Hz (Fig. 3). However, hippocampal theta and sniffing were not spectrally coherent in our data (Figs. 6 and 7). This is consistent with previous findings regarding whisking and hippocampal theta (Berg et al., 2006) and with the current consensus that theta and the exploratory rhythms originate from different oscillatory sources (Kleinfeld et al., 2016, Zhong et al., 2017). Ito et al. (2014), showed that respiration-locked olfactory bulb activity seems to be a main drive for delta oscillations and gamma power modulation in the whisker barrel cortex in the awake state. However, it has been reported that during sensory discriminations, hippocampal theta and sensory-motor rhythms can be coherent, both in the olfactory (Macrides et al., 1982; Kay, 2005) and the whisker system (Grion et al., 2016), although there are exceptions (Martin et al., 2007). Furthermore, a series of articles which have systematically explored the relation between hippocampal theta (Yanovsky et al., 2014; Lockmann et al., 2016; Nguyen Chi et al., 2016), have shown that in some conditions, especially during rest or quiescent periods, breathing activity can modulate hippocampal rhythms. The authors suggest that their results support the idea of respiration as a ‘ubiquitous channel for neural communication across the brain’ (Nguyen Chi et al., 2016).

**Figure 6.**
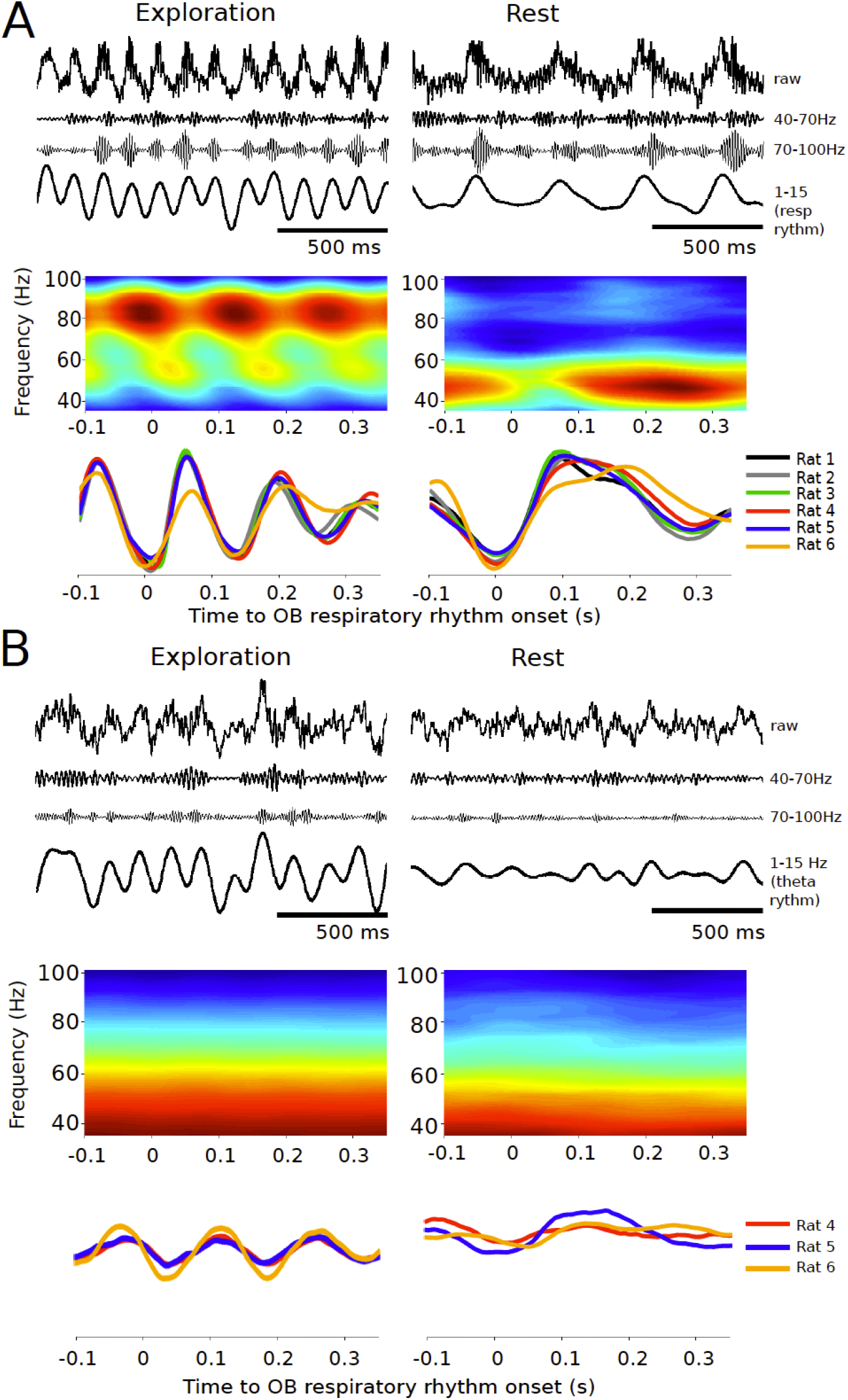
Oscillations in the olfactory bulb and the dorsal hippocampus. *A. Top*: example traces of raw and filtered OB signals, for both exploration and rest states. *Middle*: average spectrogram of OB LFP, time-locked to the onset of the OB’s respiratory rhythm. The spectrogram is an average across all respiratory rhythm cycles of a single rat. *Bottom*, mean LFP waveform, time-locked to the onset of the respiratory rhythm. Each trace represents the average of all the respiratory rhythm cycles for a different rat. *B*: Plots as in *A*, for the hippocampal signal. Example data traces in *A* are from Rat 4, and example data traces in *B* are from Rat 6.

**Figure 7.**
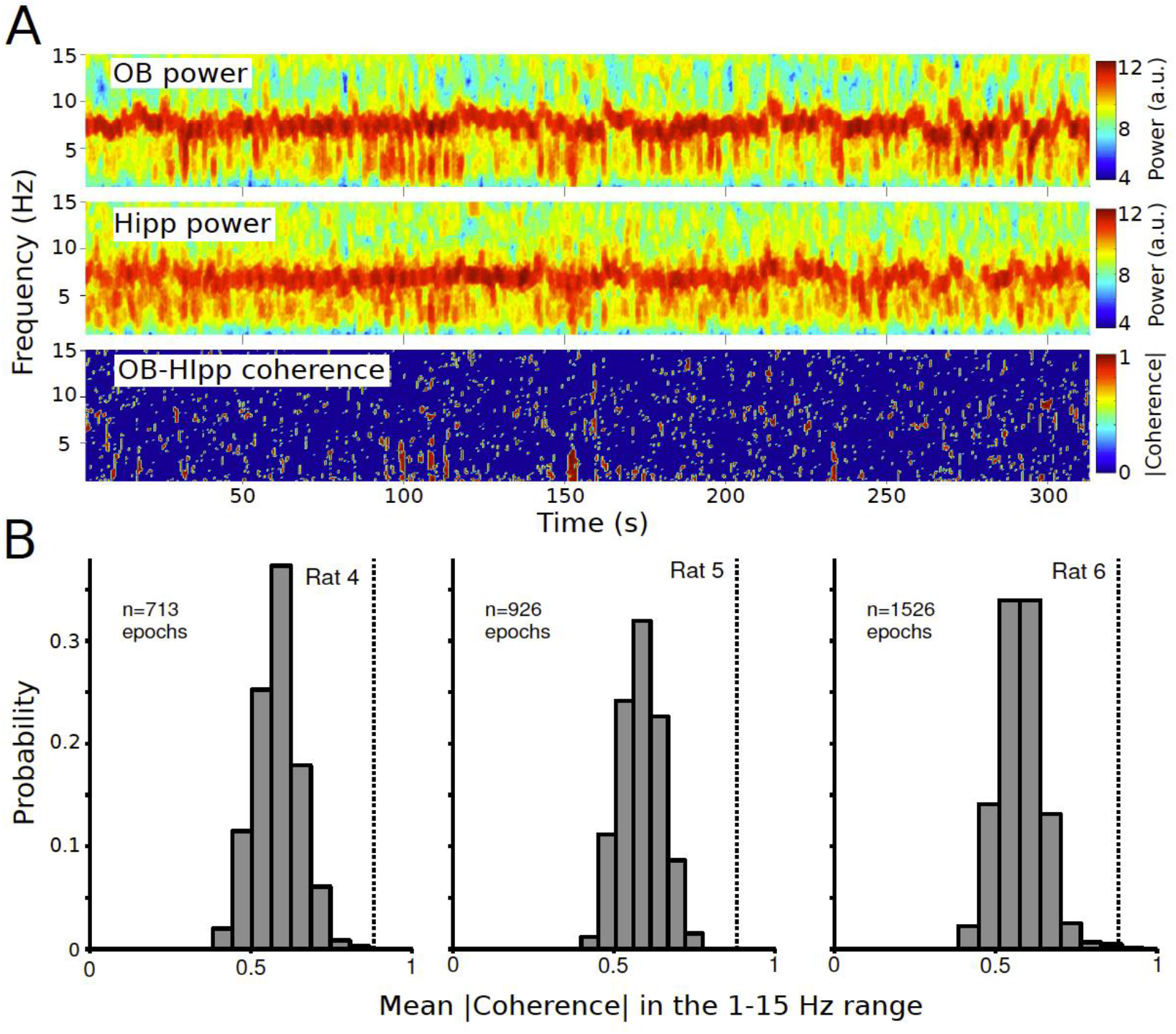
Spectral power and coherence across time for a session, for OB and hippocampal LFPs. *A. Top*: OB spectrogram. *Middle*: Hippocampal spectrogram. *Bottom*: Coherencegram. Note the common frequency range of large power (5-10 Hz) for both signals and the lack of coherence between them. For the coherence plot, only statistically coherent (*t,f*) pairs kept the actual coherence value. The rest were all set to zero (dark blue). Data are from Rat 4. *B*: Histograms estimating the probability density functions of coherence values for each rat. We computed coherence values in the 1-15 Hz for non-overlapping, 2-s epochs across all sessions and then calculated the mean absolute value of the complex-valued coherence for each epoch. We display here the probability density functions of those means, and the statistical confidence level (grey vertical lines). Coherence values larger than the confidence level are considered statistically significant. Note that in general the OB and the hippocampal time series share a frequency range but they are not spectrally coherent.

Several studies have started to show respiratory modulations across different brain areas, in both rodents and humans (Biskamp et al., 2017; Herrero et al., 2018; Ito et al., 2014; Tort et al., 2017; Zhong et al., 2017). A recent hypothesis has put together a large corpus of data regarding the breathing and whisking oscillators, proposing that the breathing rhythm would serve as a ‘master clock’ to coordinate activity from several areas (Moore et al., 2013; Kleinfeld et al., 2014). Some of these reports seem to imply that there is a difference between volitional and more automatic or basal breathing (Herrero et al., 2018) in terms of the modulation of local activity by the respiratory rhythm. Regarding neural mechanisms that could implement the reported modulations, some of them seem to depend on an intact olfactory bulb (Ito et al., 2014). In this case, the mechanical impact of the air in the nose upon each breath cycle is picked up by the olfactory sensory neurons (Grosmaitre et al., 2007), which results in the neural OB’s respiratory rhythm. This is a sort of re-entrant or ‘re-afferent’ signal, i.e., a sensory signal which results from the animal’s own motor actions (von Holst, 1954; Crapse and Sommer, 2008). The OB, in turn, projects to several forebrain areas, mainly to the pyriform and entorhinal cortices, providing a pathway for the spreading of respiratory rhythms, something which has been suggested previously (Fontanini and Bower, 2006). Another anatomical pathway that could serve as the carrier of respiratory modulations are the internal projections from the brainstem breathing oscillator to different brain areas. In this case the signal is similar to what have been described as a ‘corollary discharge’, and several pathways have been described which could perform this function (Moore et al., 2013; Yackle et al., 2017).

Altogether, our results are in line with recent reports suggesting that the timing of sensory-motor sampling rhythms can modulate the local activity of several, distant brain areas, and it can potentially function as a reference frame for the integration of activity across areas.

## ACKNOWLEDGEMENTS

We thank Cristian Lopez and Cecilia Babul for technical assistance and help with animal handling and care, and Cristóbal Cordova for help with headstage assembly. We also thank Donald Frederick for helpful comments on a previous version of the article and Francisco Parada for advice regarding data analysis.

## DISCLOSURES

The authors declare that the research was conducted without any actual or potential conflict of interest, financial or otherwise.

## Notes

**Funding:** This work was supported by: -Iniciativa Científica Milenio P10-001-F and P09-015-F -Fondo Nacional de Desarrollo Científico y Tecnológico (FONDECYT), Proyecto 3120185 (DR-L) -Beca Doctorado CONICYT (JWdS)

